# TMEM106B modifies TDP-43 pathology in human ALS brain and cell-based models of TDP-43 proteinopathy

**DOI:** 10.1101/2021.05.07.442949

**Authors:** Fei Mao, John Robinson, Travis Unger, Marijan Posavi, Defne Amado, Lauren Elman, Murray Grossman, David Wolk, Edward Lee, Vivianna M. Van Deerlin, Sílvia Porta, Virginia Lee, John Trojanowski, Alice S. Chen-Plotkin

## Abstract

The neurodegenerative diseases amyotrophic lateral sclerosis (ALS) and frontotemporal lobar degeneration with TAR DNA-binding protein-43 (TDP-43) inclusions (FTLD-TDP) share the neuropathological hallmark of aggregates of TDP-43. However, factors governing the severity and regional distribution of TDP-43 pathology, which may account for the divergent clinical presentations of ALS and FTLD-TDP, are not well-understood. Here, we investigated the influence of genotypes at *TMEM106B*, a locus associated with risk for FTLD-TDP, and hexanucleotide repeat expansions in *C9orf72*, a known genetic cause for both ALS and FTLD-TDP, on global TDP-43 pathology and regional distribution of TDP-43 pathology in 899 postmortem cases from a spectrum of neurodegenerative diseases. We found that, among the 110 ALS cases, minor (C)- allele homozygotes at the *TMEM106B* locus sentinel SNP rs1990622 had more TDP-43 pathology globally, as well as in select brain regions. *C9orf72* expansions similarly associated with greater TDP-43 pathology in ALS. However, adjusting for *C9orf72* expansion status did not affect the relationship between *TMEM106B* genotype and TDP-43 pathology. In order to elucidate the direction of causality for this association, we directly manipulated *TMEM106B* levels in an inducible cell system that expresses mislocalized TDP-43 protein. We found that partial knockdown of *TMEM106B*, to levels similar to what would be expected in rs1990622 C allele carriers, led to development of more TDP-43 cytoplasmic aggregates, which were more insoluble, in this system. Taken together, our results support a causal role for TMEM106B in modifying the development of TDP-43 proteinopathy.

## INTRODUCTION

Neurodegenerative diseases are a major and growing cause of morbidity and mortality worldwide, especially in the elderly. Progress towards disease-modifying therapies has been slow, largely owing to an incomplete understanding of disease pathogenesis. Abnormal protein aggregates in stereotypical brain regions characterize the various neurodegenerative diseases neuropathologically, with prototypical examples including tau-positive neurofibrillary tangles and amyloid-β-positive plaques in Alzheimer’s disease (AD), α-synuclein aggregates in Lewy body disease (LBD), and TAR DNA-binding protein 43 (TDP-43)-positive inclusions in amyotrophic lateral sclerosis (ALS) and frontotemporal lobar degeneration with TDP-43 inclusions (FTLD-TDP). However, overlap exists among the neurodegenerative diseases, as exemplified by multiple co-occurring proteinopathies in approximately half of neurodegenerative disease cases [17].

TDP-43 was first identified in 2006 as the main protein component of the ubiquitinated inclusions found in ALS and FTLD-U [14], which was subsequently termed FTLD-TDP [12]. The discovery of this shared “proteinopathy,” along with subsequent reports of rare genetic mutations – *e.g.* in *TARDBP*, the gene that encodes TDP-43 – that could cause both ALS and FTLD-TDP, has led to our current understanding of these two disease presentations as ends of a shared pathophysiological spectrum [3]. Indeed, since first reports in 2011 [6,16], a hexanucleotide (GGGGCC) repeat expansion in the *C9orf72* gene has proven to be the most frequent genetic cause of both ALS and FTLD-TDP. Moreover, also in 2011, we reported that common variants at the *TMEM106B* locus found by genome-wide association to increase risk for developing FTLD-TDP also associated with risk for developing cognitive impairment in ALS [23,24].

Despite these commonalities and the presence of overlapping syndromes, clinical presentation and the pattern of TDP-43 inclusions do show some differences in ALS and FTLD-TDP. In particular, pathological lesions and neurodegeneration mainly affect the upper and lower motor neurons in ALS while predominantly affecting the frontal and temporal lobes in FTLD-TDP. Clinical symptomatology accordingly follows from the functional neuroanatomy of the most involved areas of the central nervous system (CNS).

What factors influence, then, where and to what extent TDP-43 aggregates may form in the CNS? To answer this question, we assessed the pattern of TDP-43 deposition in 899 autopsied individuals representing a broad spectrum of clinical neurodegenerative diseases, focusing on the influence of *TMEM106B* genotype and *C9orf72* expansion on the regional distribution and global burden of TDP-43 pathology. Moreover, to better understand whether *TMEM106B* genotype effects might be *causal* for modifying the development of TDP-43 pathology, we directly manipulated *TMEM106B* levels in a cell-based model.

## MATERIALS AND METHODS

### Cohorts

899 patients (Table 1) with a neuropathological diagnosis of neurodegenerative disease (ALS, FTLD-TDP, AD or LBD) were identified within a previously-described autopsy database at the University of Pennsylvania (Penn) Center for Neurodegenerative Disease Research (CNDR) [26]. Demographic, clinical, and neuropathological data were obtained for all autopsy cases conducted at the CNDR from 1989 to 2019. Informed consent was obtained for all participants prior to death. All procedures in these studies adhere to the tenets of the Declaration of Helsinki.

**Table 1.** Demographic characteristics

|  | ALS | FTLD-TDP | AD | LBD |
| --- | --- | --- | --- | --- |
| *n* = | 110 | 99 | 523 | 167 |
| Sex, % |  |  |  |  |
| Male | 66 (60%) | 55 (56%) | 238 (46%) | 132 (79%) |
| Female | 44 (40%) | 44 (44%) | 285 (54%) | 35 (21%) |
| Disease duration, yr | 3 (2-5) | 7 (5-10) | 9 (6-12) | 12 (9-18) |
| Age at death, yr | 63 (56-71) | 70 (63-75) | 79 (70-85) | 79 (72-83) |

Disease duration and Age at death are shown as median (IQRs).

### Cognitive testing

In the ALS cohort, antemortem cognitive testing was performed at 3–6-month intervals during routine clinic visits. Data for a test of letter-guided verbal fluency (FAS test [11]) was available for 52/110 patients (median time between test and death = 2 years, IQR 1-2 years). For the FAS test, patients without significant dysarthria were asked to say as many unique words beginning with the target letter “F” as possible in one minute. Patients with significant dysarthria but preserved limb and hand function were asked to write the words; 90 seconds was allotted to these patients. The total number of unique words produced was recorded. We have previously shown that both tests perform well compared to the Frontal Behavioral Inventory (FBI), a battery of neuropsychological tests used in assessing cognitive impairment in FTLD. On receiver operating curve analysis compared to the FBI, the FAS test has an area under the curve (AUC) of 0.88 in differentiating cognitively intact and cognitively impaired individuals.

### Neuropathological staging

20 CNS regions (Table 2) encompassing the brain and spinal cord were examined in the CNDR neuropathology evaluations as previously described [14]. Briefly, each region was assigned a semiquantitative score (0 = none, 0.5 = rare, 1 = mild, 2 = moderate, 3 = severe) for TDP-43 pathology, based on immunohistochemical staining (1D3 antibody for phosphorylated TDP-43, provided by Dr. M Neumann and Dr. E. Kremmer) without prior knowledge of the clinical diagnosis.

**Table 2.** Severity of TDP-43 pathology in each region in ALS cases

| TDP-43 pathology scores |  |  |
| --- | --- | --- |
|  | Median | IQRs |
| 01 Spinal cord | 2 | 2-3 |
| 02 Motor cortex | 2 | 0.5-2 |
| 03 Medulla | 1 | 0.625-2 |
| 04 Caudate/Putamen | 1 | 0-1 |
| 05 Midbrain | 0.5 | 0-1 |
| 06 Globus Pallidus | 0.5 | 0-1 |
| 07 Anterior cingulate | 0.5 | 0-1 |
| 08 Thalamus/subthalamus | 0.5 | 0-1 |
| 09 Pons | 0.5 | 0-1 |
| 10 Substantia nigra | 0.5 | 0-1 |
| 11 Middle frontal cortex | 0 | 0-1 |
| 12 Amygdala | 0 | 0-0.75 |
| 13 Angular gyrus | 0 | 0-0.5 |
| 14 CA1/subiculum | 0 | 0-0.5 |
| 15 Entorhinal cortex | 0 | 0-0.5 |
| 16 Superior/middle temporal cortex | 0 | 0-0.5 |
| 17 Dentate gyrus | 0 | 0-0.375 |
| 18 Cerebellum | 0 | 0-0 |
| 19 Locus coeruleus | 0 | 0-0 |
| 20 Occipital cortex | 0 | 0-0 |

### Genotyping

DNA was extracted from brain samples, and genotyping was performed using Taq-Man chemistry-based allelic discrimination assays (Applied Biosystems, Foster City, CA, USA) as previously described [23]. The sentinel SNP rs1990622 at the *TMEM106B* locus was genotyped. Genetic analysis of *C9orf72* for hexanucleotide repeat expansions was performed as previously described [19]. Briefly, PCR products were separated by capillary electrophoresis on an ABI PRISM 3130xl Genetic Analyzer (Applied Biosystems) followed by analysis with GeneMapper software v3.7. A repeat size of greater than or equal to 30 repeats was used as the cut-off for an expansion.

### Genetic ancestry analysis

To assess genetic ancestry, we began with a panel of ancestral informative markers (AIMs) generated and validated by Kosoy *et al.*, with previously-demonstrated ability to discern individuals from different ancestry groups using as few as 24 markers [13]. ALS patients were genotyped for AIMs on the Illumina Infinium Global Screening Array (63 ALS individuals genotyped for 88 AIMs) and on Illumina OmniExpress array (43 ALS individuals genotyped for 82 AIMs). Merging the two datasets, we extracted 63 AIMs present in both datasets, creating a final dataset of 63 AIM genotypes for 106 out of 110 ALS individuals. AIM genotypes were not available for the remaining four ALS patients, and they were excluded from the analysis.

We clustered the 106 ALS individuals by performing principal components analysis (PCA) on genotypes of these 63 AIMs. To obtain PCA coordinates we employed *prcomp* [20], while for visualization we used *fviz_pca_ind* R function [10]. We performed outlier detection using squared Mahalonobis distances (MD) to account for data correlations and multidimensionality of the data. We estimated MD using the orthogonalized Gnanadesikan-Kettenring pairwise method. For this purpose we employed the R function *covRob* with *pairwiseGK* estimator [25]. We obtained the outlier p-values using cumulative chi-squared distribution and used the Bonferroni correction to adjust the p-values for multiple hypothesis testing.

### Cell culture, transfection and doxycycline induction

A doxycycline (Dox)-inducible green fluorescent protein (GFP)-tagged cytoplasmic TDP-43 stable cell line (iGFP-NLSm) was previously established by Sílvia Porta *et al.*[15] in QBI-293 cells. Briefly, upon addition of doxycycline to the media, transgenic expression of a form of TDP-43 with mutations introduced in the nuclear localization signal (NLS) is achieved, with subsequent localization of GFP-TDP-43NLSm protein in the cytoplasmic compartment. For *TMEM106B* or control knockdown, we employed transient transfection with previously-validated shRNA constructs, using Lipofectamine 2000 (Invitrogen) 1 day after plating (80-90% confluency) according to manufacturer instructions under serum-free conditions. Transfection medium was then replaced by whole growth medium with or without doxycycline (1μg/mL).

### TDP-43 Immunofluorescence

iGFP-NLSm stable cells were plated on poly-D-lysine coated coverslips. Cells were washed once with dPBS at room temperature, fixed 15 min at room temperature with 4% paraformaldehyde (4% PFA) containing 1% Triton X-100 to remove soluble proteins, then washed three times with dPBS. After incubation in blocking buffer (3% bovine serum albumin–3% FBS in dPBS) for 30 min at room temperature, cells were incubated with specific primary antibodies overnight at 4 °C in a humidified chamber. After washing with dPBS, cells were incubated with appropriate secondary antibodies in blocking buffer for 1 hour at room temperature in the dark. Coverslips were mounted on slides (ProLong Gold, Invitrogen).

The following antibodies and conditions were used: anti-TDP-43: TDP-43 C1038 antibody (gift of V.M. Lee and J. Trojanowski, CNDR) was used at 0.3375 μg/mL (1:10,000 dilution) [5]; anti-GFP antibody (catalog # MAB2510, Millipore) was used at 1 μg/mL. All secondary antibodies were Alexa Fluor antibodies (Invitrogen) used at 2 μg/mL.

### Lysosome characterization

iGFP-NLSm stable cells were plated onto coverslips, transfected with shTmem106b or control, and induced with doxycycline as described above. 72 hours after transfection, cells were treated with Lysotracker 594 (Invitrogen) at a dilution of 1:1000 in standard media and incubated at 37 degrees C for 30 minutes. After this incubation, cells were fixed for fifteen minutes in 2% paraformaldehyde, washed 4 times with PBS and then incubated with Draq5 (Invitrogen) for 30 minutes. The coverslips were then placed onto slides using mounting media (ProLong Gold, Invitrogen). Lamp1 was stained by fixing cells using 2% paraformaldehyde, washing with PBS, and blocking using 3% BSA, 0.05% saponin for one hour. Lamp1 antibody H4A3 (DSHB) was diluted into the blocking buffer at 1 μg/mL and allowed to incubate overnight at 4 degrees C in a humid chamber. Alexa 594 (Invitrogen) secondary was used at a dilution of 1:1000 and incubated for one hour. Cells were then washed and incubated with Draq5 for 30 minutes, then mounted onto slides (ProLong Gold, Invitrogen). Images were captured using a Leica SP5 confocal microscope.

### Protein preparation and immunoblotting

Sequential biochemical fractionation of cell lysates was performed. Cells were washed with dPBS and extracted with RIPA buffer (50 mM Tris, 150 mM NaCl, 5 mM EDTA, 0.5% sodium deoxycholate, 1% NP-40, 0.1% SDS, pH 8.0) and then centrifuged at 21,380 × *g* for 30 min at 4□. The supernatants were saved for analysis as the soluble fraction, and pellets were re-suspended with 2% SDS. After a second round of centrifugation at 21,380 × *g* for 30 min at 4□, the supernatants were collected as the RIPA-insoluble fraction. Both RIPA-soluble and RIPA-insoluble fractions were prepared for immunoblotting. The following antibodies and conditions were used: anti-TMEM106B: TMEM106B 3650 antibody (rabbit polyclonal antibody raised in-house against amino acids 36-50) was used at 1:500 dilution, anti-TDP-43 antibody (catalog # 10782-2-AP, Proteintech) was used at 0.307 μg/mL (1:1000 dilution); anti-GAPDH antibody (catalog # 2-RGM2, Advanced ImmunoChemical) was used at 1 μg/mL.

### Statistical Analysis

Linear regression analyses were used to evaluate the association of factors (*TMEM106B* genotype, *C9orf72* expansion, age at death) with global or regional TDP-43 pathology scores. Kruskal-Wallis rank sum tests were performed to make comparisons of TDP-43 pathology score between disease subgroups. For cell-based experiments, one-way ANOVAs were used to compare *TMEM106B* knockdown with control (scrambled shRNA) conditions, followed by Tukey’s posthoc testing for pairwise comparisons, with correction for multiple comparisons. All statistical analysis was performed in R (http://www.r-project.org). The maps used to illustrate the distribution of TDP-43 pathology and coefficients were created using BioRender (https://biorender.com/).

## RESULTS

### Cohort characteristics

We examined a total of 899 cases from an autopsy database of neurodegenerative disease cases collected at the University of Pennsylvania (Penn) between 1989 and 2019. Primary neuropathological diagnoses were ALS (*n*=110), FTLD-TDP (*n*=99), AD (*n*=523), and LBD (*n*=167). Demographic and clinical data for these cases are shown in Table 1.

For each CNS region (Table 2, *n*=20 regions: amygdala, angular gyrus, cerebellum, anterior cingulate, caudate/putamen, CA1/subiculum, dentate gyrus, entorhinal cortex, globus pallidus, locus coeruleus, motor cortex, midbrain, medulla, middle frontal cortex, occipital cortex, pons, spinal cord, superior/middle temporal cortex, substantia nigra and thalamus/subthalamus), the extent of TDP-43 pathology was assigned a semiquantitative score (0=none, 0.5=rare, 1=mild, 2=moderate, and 3=severe, Fig. 1 a-d), as previously described [21], and these scores were then averaged across all CNS regions to obtain a global TDP-43 score for each case. FTLD-TDP cases had the highest global TDP-43 scores, with a median score of 1.66 (IQR = 1.41-1.83), followed by ALS cases (median score 0.74, IQR 0.37-1.00). AD and LBD cases showed low levels of TDP-43 pathology (Supplementary Fig. 1a).

**Fig. 1.**
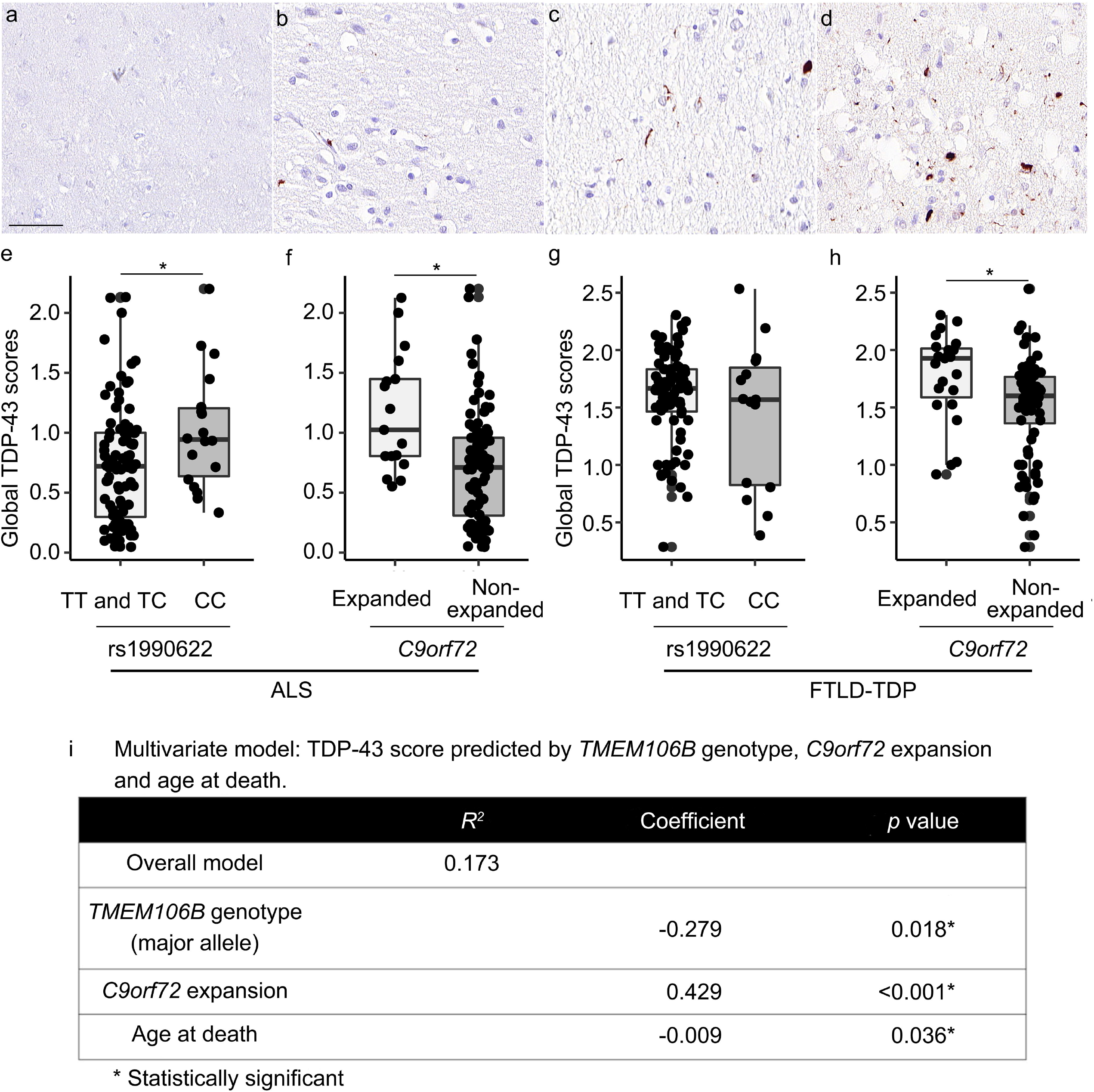
TDP-43 immunohistochemistry in anterior cingulate brain sections illustrate the spectrum of pathology (**a**-**d**). Each region assessed was assigned a semiquantitative score for TDP-43 pathology: 0 = none (**a**), 0.5 = rare, 1 = mild (**b**), 2 = moderate (**c**) or 3 = severe (**d**). “Rare” is defined as a mild level of TDP-43 pathology in a single field of view. Global TDP-43 pathology scores differ significantly among *TMEM106B* genotypes under a major (T)-allele dominant model in ALS (**e**) but not in FTLD-TDP (**g**). *C9orf72* expansion significantly associates with the global extent of TDP-43 pathology in both ALS (**f**) and FTLD-TDP (**h**). Scale bar = 50 μm.

*TMEM106B* genotypes at the sentinel SNP rs1990622 are shown in Table 3, along with the frequency of *C9orf72* expansions found in each disease group. Notably, 15% of our ALS cohort and 23% of our FTLD-TDP cohort harbored *C9orf72* repeat expansions.

**Table 3.** *TMEM106B* rs1990622 genotype and *C9orf72* expansion frequencies among neurodegenerative disease groups

|  |  | ALS | FTLD-TDP | AD | LBD |
| --- | --- | --- | --- | --- | --- |
| *TMEM106B*<br>rs1990622 | TT | 38 (35%) | 44 (44%) | 149 (28%) | 47 (28%) |
|  | TC | 54 (49%) | 40 (40%) | 254 (49%) | 88 (53%) |
|  | CC | 18 (16%) | 15 (15%) | 120 (23%) | 32 (19%) |
| *C9orf72* | Expanded | 17 (15%) | 23 (23%) | 1 (0%) | 0 (0%) |
|  | Non-expanded | 93 (85%) | 76 (77%) | 522 (100%) | 167 (100%) |

All values are shown as *n* (% of disease group).

### *TMEM106B* genotype and presence of *C9orf72* expansion both associate with global extent of TDP-43 pathology

Within each disease group, we next asked whether *TMEM106B* rs1990622 genotype modified global TDP-43 pathology scores. We employed both codominant and major (T)-allele-dominant models, given existing literature suggesting that minor (C)- allele homozygotes differ from both major (T)-allele homozygotes and heterozygotes [9,23]. For the 110 ALS cases, global TDP-43 pathology scores differed significantly among *TMEM106B* genotypes under a major (T)-allele dominant model (*p* = 0.018), with homozygotes for the minor allele (CC) having the highest global TDP-43 pathology scores (Fig. 1e, Supplementary Fig. 2). Global TDP-43 scores did not differ among carriers of different *TMEM106B* genotypes in FTLD-TDP cases (Fig. 1g), AD cases (Supplementary Fig. 1b), or LBD cases (Supplementary Fig. 1c). As there was relatively little TDP-43 pathology in AD cases and LBD cases, we additionally analysed the subset of AD and LBD cases with the most abundant TDP-43 pathology (highest quartile of TDP-43 pathology score). No significant *TMEM106B* genotype effects were found in this subset analysis (Supplementary Fig. 1d and 1e).

We performed similar analyses comparing global TDP-43 pathology scores in individuals with vs. without *C9orf72* expansions. In both ALS and FTLD-TDP, *C9orf72* expansion carriers showed higher global TDP-43 pathology scores (*p* = 0.001 for ALS, *p* = 0.009 for FTLD-TDP, Fig. 1f and 1h). There were no individuals with *C9orf72* expansions in the LBD group and only one individual with *C9orf72* expansion in the AD group.

We assessed whether clinical covariates such as age at death, sex, or total disease duration might also impact global TDP-43 pathology scores, finding that only age at death was significantly associated with global TDP-43 pathology scores (β = −0.011, *p* = 0.015 for ALS; β = −0.015, *p* = 0.001 for FTLD-TDP).

Given the significant individual effects of age at death, *TMEM106B* genotype, and *C9orf72* expansion on global TDP-43 pathology scores in ALS, we tested these three variables together in a linear regression model predicting global TDP-43 pathology score in ALS cases. Co-varying for the other two variables did not remove significant associations for any of these variables (*R*^2^ = 0.173 for full model; *β* = −0.009, *p* = 0.036 for age at death; *β* = −0.278, *p* = 0.018 for *TMEM106B* genotype; *β* = 0.429, *p* < 0.001 for *C9orf72* expansion).

Finally, because genetic ancestry may affect results, we performed a genetic ancestry analysis using 63 ancestry informative markers (AIMs, Supplementary Table 1) in 106/110 individuals from the ALS cohort for which we could obtain further genotype data. As shown in Supplementary Fig. 3, five individuals demonstrated substantially different genetic ancestry based on principal components analysis (PCA). We repeated our analyses excluding these five individuals as well as the four individuals for whom we could not ascertain AIMs. Neither the association between *TMEM106B* genotype and global TDP-43 pathology scores, nor the association between *C9orf72* expansion status and global TDP-43 pathology scores were substantially affected (Supplementary Fig. 4).

### The extent of TDP-43 pathology in specific brain regions is impacted by *TMEM106B* genotype

Having demonstrated a relationship between *TMEM106B* genotype and global TDP-43 pathology score in ALS, we sought to further characterize this relationship in individual CNS regions. As expected, regions with the most severe TDP-43 pathology in the ALS cases were the spinal cord and motor cortex (Table 2, Fig. 2a-b). However, less severe TDP-43 pathology was found in many other regions (Table 2, Fig. 2a-e).

**Fig. 2.**
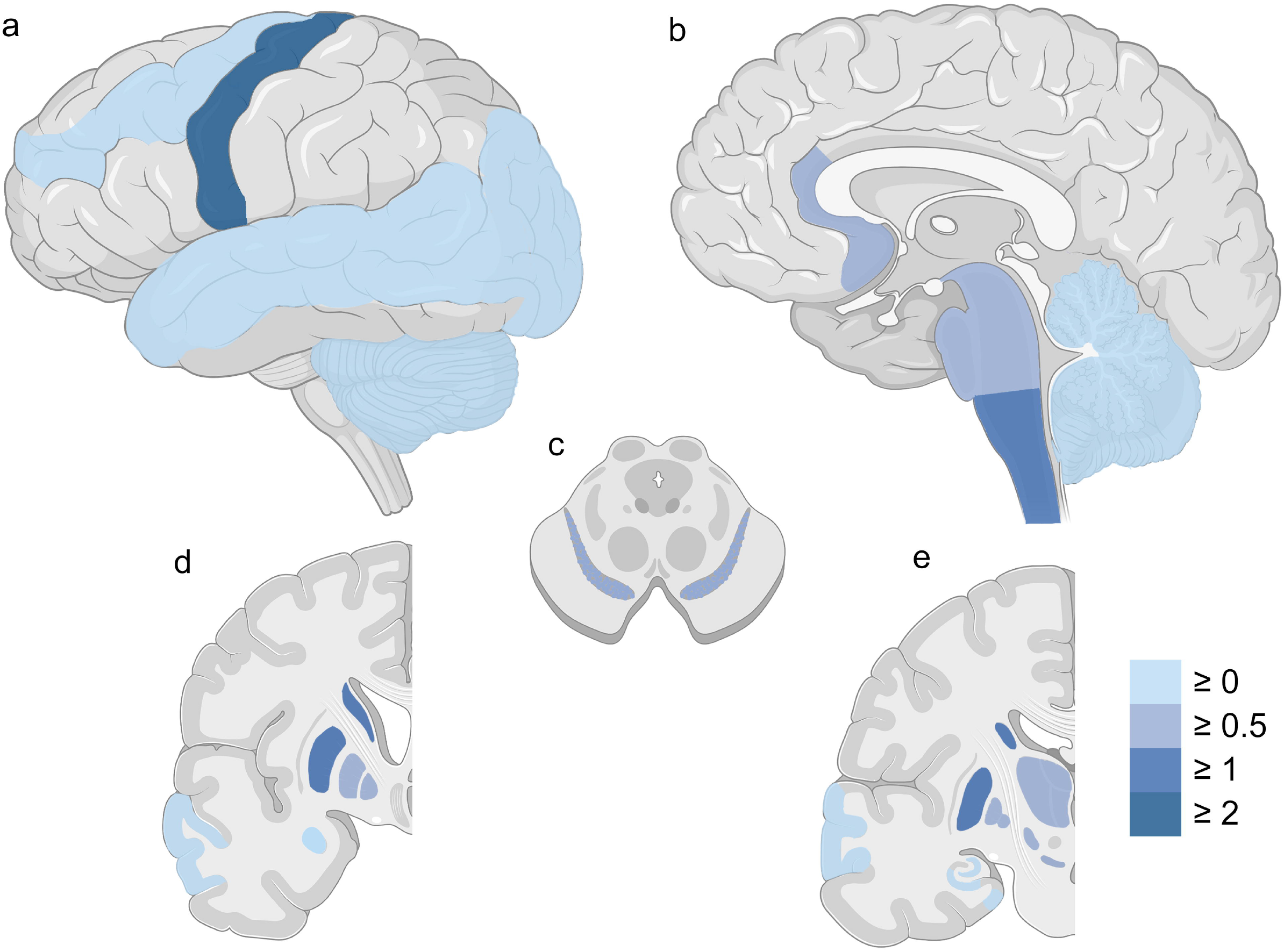
Severity of TDP-43 pathology in different brain regions in ALS cases. Lateral (**a**), sagittal (**b**), cross-sectional (**c**), and coronal (**d**, **e**) sections of CNS regions are shown. Colors reflect the calculated median severity scores across all involved cases. The color scale depicts median TDP-43 pathology from none (0 = light blue) to moderate or severe (>2 = dark blue) in each region. Spinal cord is not shown.

We thus asked whether *TMEM106B* genotype associated with TDP-43 pathology score in each of these CNS regions in the ALS cohort. Seven regions showed nominally-significant associations between severity of TDP-43 pathology and *TMEM106B* genotype under a major (T)-allele-dominant model: amygdala (*p* = 0.028), angular gyrus (*p* = 0.026), anterior cingulate (*p* = 0.023), midbrain (*p* = 0.017), middle frontal cortex (*p* = 0.022), superior/middle temporal cortex (*p* = 0.011), substantia nigra (*p* = 0.004). In all seven regions, major (T) allele carriers had lower TDP-43 pathology scores (Table 4). Notably, the CNS regions in which *TMEM106B* genotype exerted the largest effects were not, in general, areas with abundant TDP-43 pathology (Fig. 3).

**Table 4.**
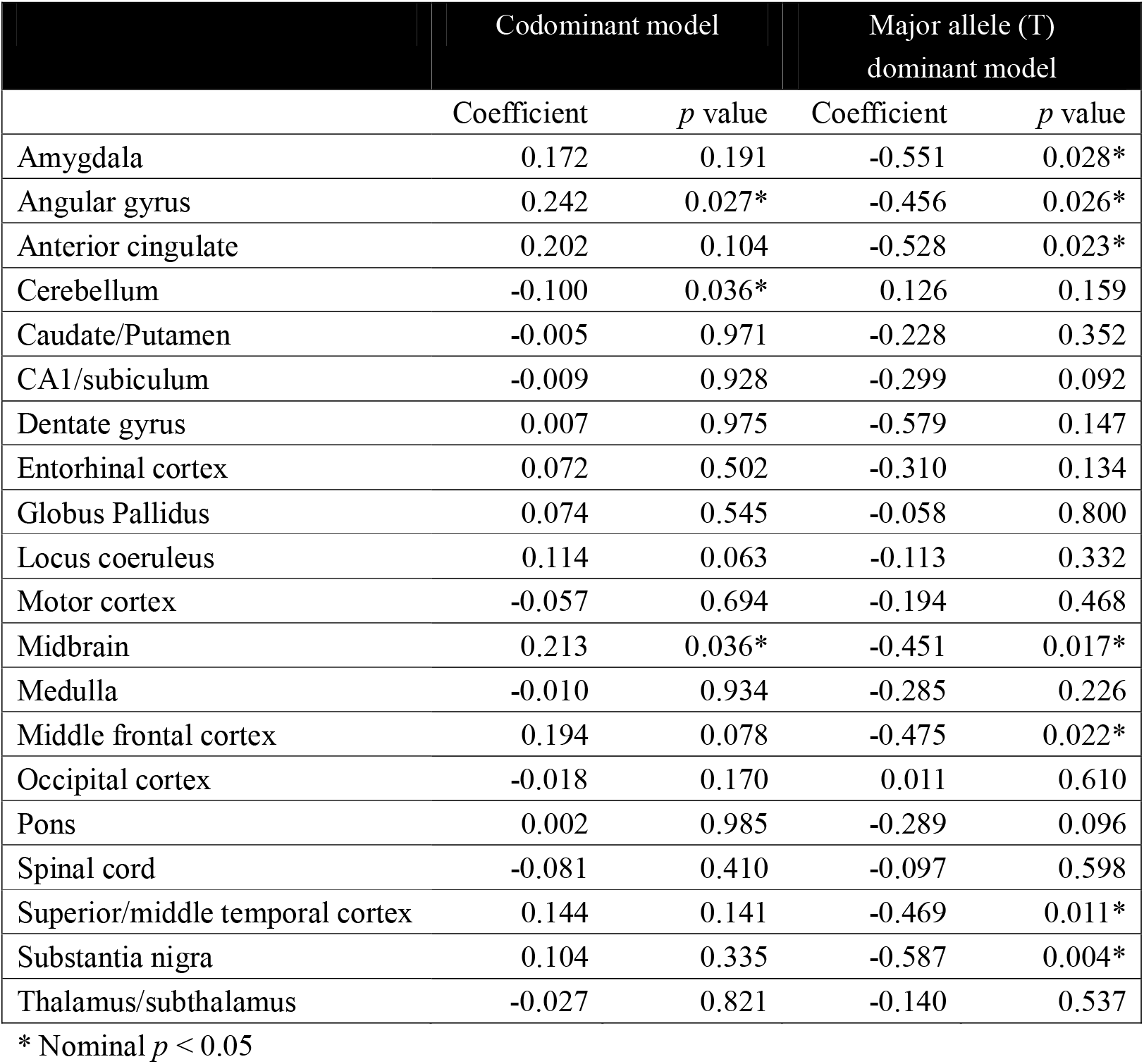
Bivariate linear model: TDP-43 scores predicted by *TMEM106B* genotype

**Fig. 3.**
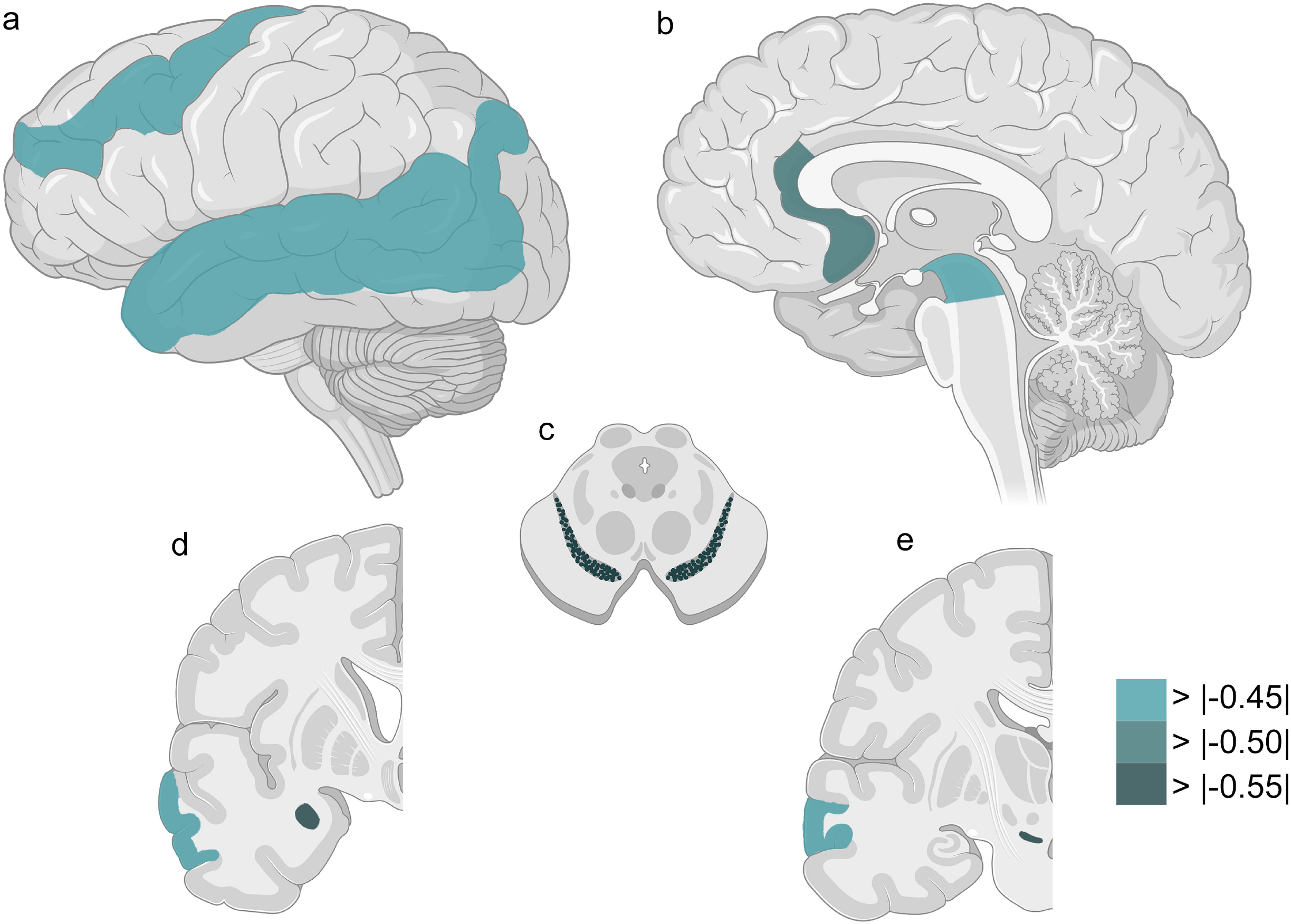
Seven brain regions show nominally-significant associations between severity of TDP-43 pathology and *TMEM106B* genotype under a major (T)-allele-dominant model in ALS. Lateral (**a**), sagittal (**b**), cross-sectional (**c**), and coronal (**d**, **e**) sections of CNS regions are shown. Colors reflect the effect size (calculated coefficients) for *TMEM106B* genotype on TDP-43 pathology across all ALS cases, ranging from weak effects (light green) to strong effects (dark green). See Table 4 for additional details.

Despite a lack of correlation between *TMEM106B* genotype and global TDP-43 pathology in FTLD-TDP cases, we performed a similar analysis to understand effects on regional deposition of TDP-43 pathology in FTLD-TDP. Only two regions showed nominally-significant associations between TDP-43 pathology score and *TMEM106B* genotype under a major (T)-allele-dominant model: caudate/putamen (*p* = 0.038), middle frontal cortex (*p* = 0.013). Of note, *TMEM106B* genotype associated with the degree of TDP-43 pathology in the middle frontal cortex in both ALS and FTLD-TDP cases, but in opposite directions (*β* = −0.475 for ALS, *β* = 0.565 for FTLD-TDP, Supplementary Table 2).

### The extent of TDP-43 pathology in specific brain regions is impacted by *C9orf72* expansion

We performed a similar analysis to understand *C9orf72* expansion effects on regional deposition of TDP-43 pathology in ALS cases. Twelve regions showed nominally-significant associations between severity of TDP-43 pathology and presence of a *C9orf72* expansion: amygdala (p = 0.002), angular gyrus (p = 0.022), CA1/subiculum (p < 0.001), dentate gyrus (p < 0.001), entorhinal cortex (p = 0.030), globus pallidus (p = 0.048), motor cortex (p = 0.015), middle frontal cortex (p = 0.014), occipital cortex (p = 0.003), pons (p = 0.010), substantia nigra (p = 0.007), thalamus/subthalamus (p = 0.006) (Table 5). Of note, the CNS regions in which *C9orf72* expansion and *TMEM106B* genotypes associated with TDP-43 pathology did not overlap substantially (Fig. 4).

**Table 5.**
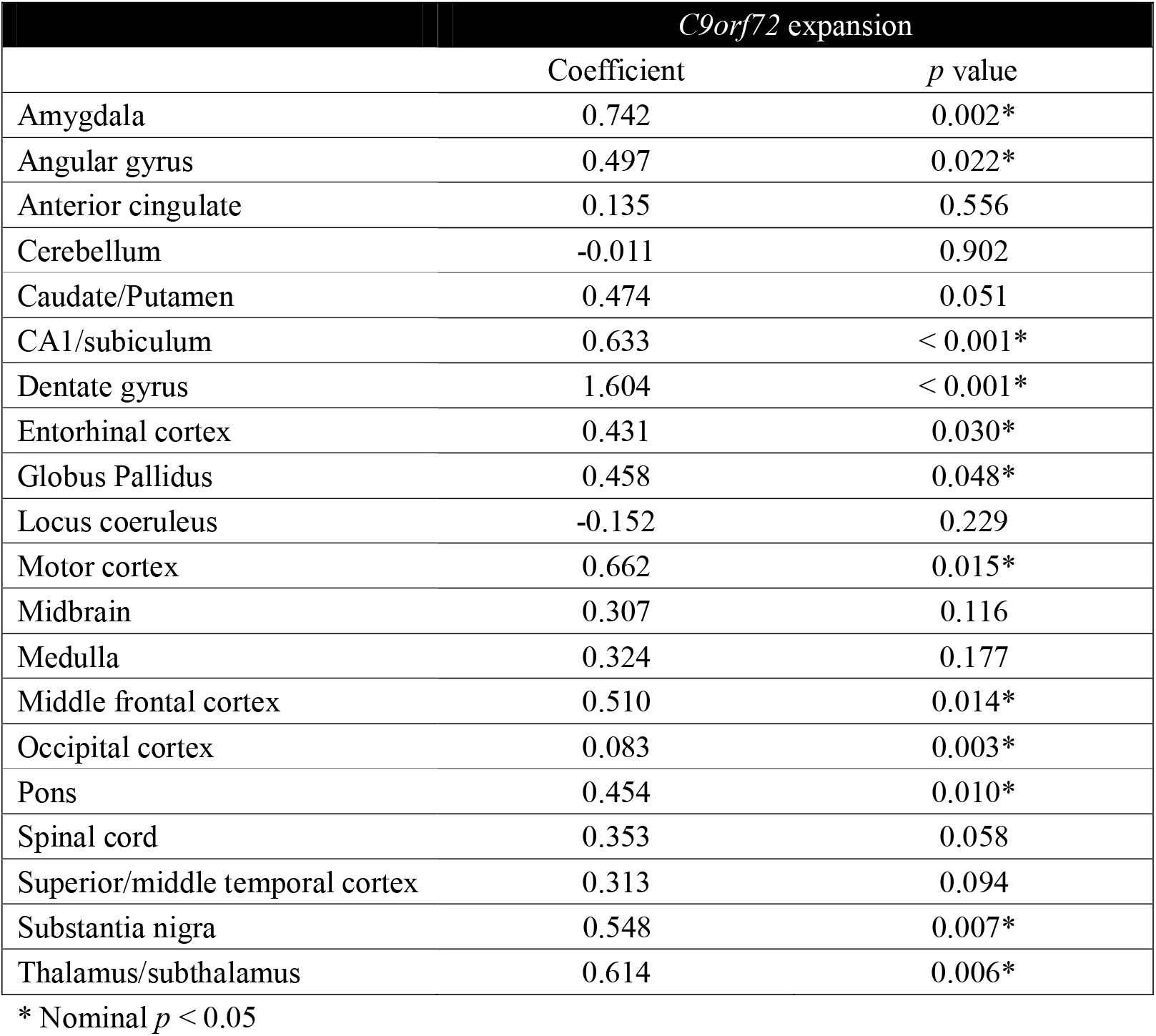
Bivariate linear model: TDP-43 scores predicted by *C9orf72* expansion

|  | *C9orf72* expansion |  |
| --- | --- | --- |
|  | Coefficient | *p* value |
| Amygdala | 0.742 | 0.002* |
| Angular gyrus | 0.497 | 0.022* |
| Anterior cingulate | 0.135 | 0.556 |
| Cerebellum | -0.011 | 0.902 |
| Caudate/Putamen | 0.474 | 0.051 |
| CA1/subiculum | 0.633 | < 0.001* |
| Dentate gyrus | 1.604 | < 0.001* |
| Entorhinal cortex | 0.431 | 0.030* |
| Globus Pallidus | 0.458 | 0.048* |
| Locus coeruleus | -0.152 | 0.229 |
| Motor cortex | 0.662 | 0.015* |
| Midbrain | 0.307 | 0.116 |
| Medulla | 0.324 | 0.177 |
| Middle frontal cortex | 0.510 | 0.014* |
| Occipital cortex | 0.083 | 0.003* |
| Pons | 0.454 | 0.010* |
| Spinal cord | 0.353 | 0.058 |
| Superior/middle temporal cortex | 0.313 | 0.094 |
| Substantia nigra | 0.548 | 0.007* |
| Thalamus/subthalamus | 0.614 | 0.006* |

\* Nominal $p < 0.05$

**Fig. 4.**
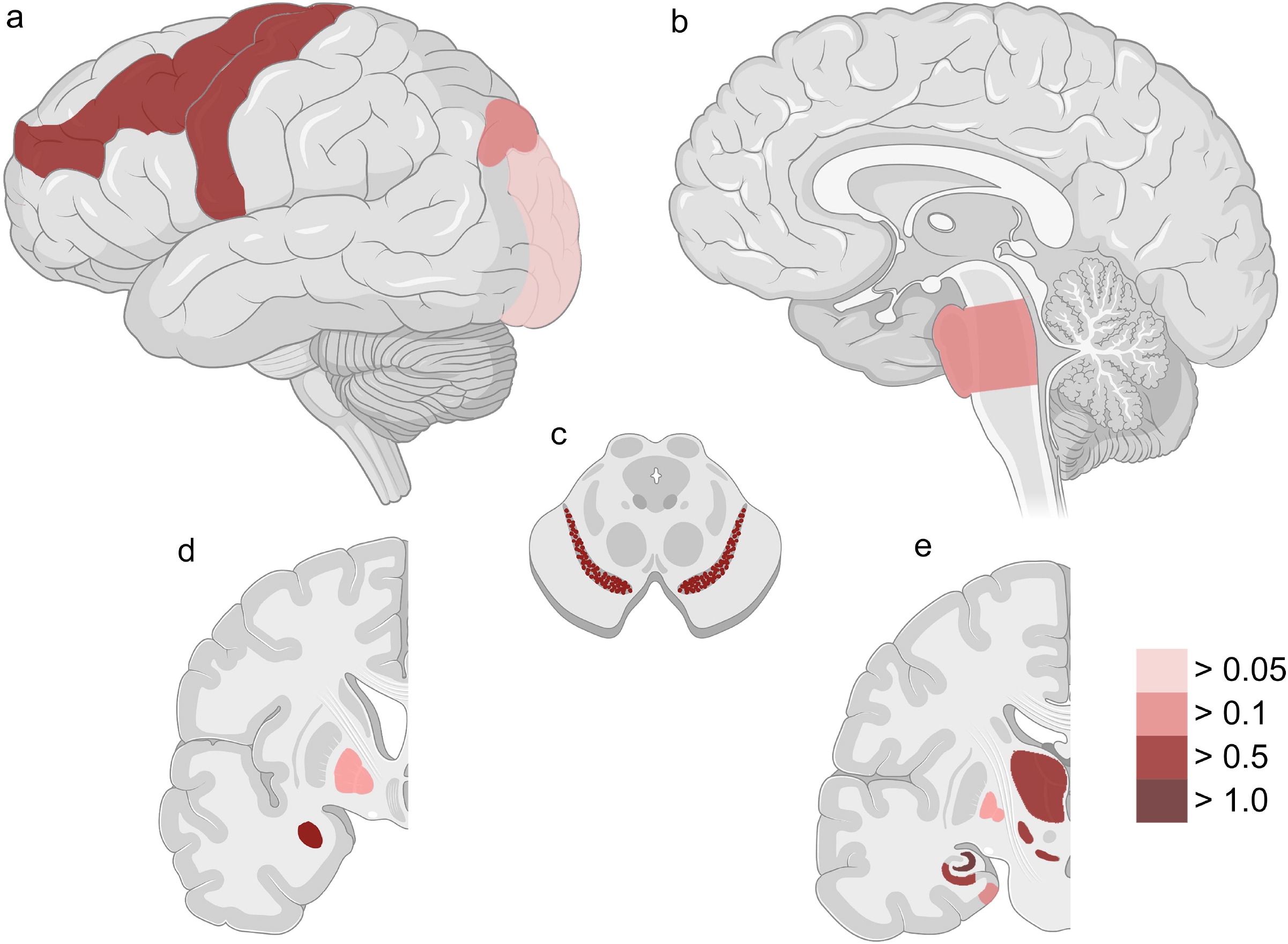
Twelve CNS regions showed significant associations between severity of TDP-43 pathology and *C9orf72* expansion in ALS. Lateral (**a**), sagittal (**b**), cross-sectional (**c**), and coronal (**d**, **e**) sections of CNS regions are shown. Colors reflect the effect size (calculated coefficients) for presence of a *C9orf72* expansion on TDP-43 pathology across all ALS cases, ranging from weak effects (light red) to strong effects (dark red). Spinal cord is not shown. See Table 5 for additional details.

We next examined FTLD-TDP cases to understand *C9orf72* expansion effects on regional deposition of TDP-43 pathology. *C9orf72* expansion modified the extent of TDP-43 pathology found in four regions in FTLD-TDP cases (dentate gyrus, midbrain, middle frontal cortex and substantia nigra), all in the same direction as for ALS (Supplementary Table 3).

### *TMEM106B* knockdown leads to increased TDP-43 proteinopathy in a cell-based system

We have previously reported that *TMEM106B* rs1990622 genotypes associate with *TMEM106B* mRNA expression levels in multiple tissue types. In lymphoblastoid cell lines, the rs1990622 major (T) allele corresponds to higher levels of *TMEM106B* expression, through alterations in chromatin architecture secondary to differential binding of CTCF to alleles at a causal variant linked to rs1990622 [8]. Because neuropathological studies do not allow us to address the question of causality, we turned next to an inducible cell system that expresses mislocalized TDP-43 protein in which we could directly manipulate levels of *TMEM106B* expression.

We employed a previously-characterized cell line expressing GFP-tagged TDP-43 (GFP-NLS-TDP) that can be targeted to the cytoplasm under doxycycline induction [15]. In this model, TDP-43 occasionally forms insoluble cytoplasmic aggregates, particularly under conditions where cells have been stressed or “seeded” with extracts from FTLD-TDP brain [15]. By manipulating *TMEM106B* expression in this cell model, we could test the hypothesis that decreased *TMEM106B* expression (associated with minor (C) allele homozygosity at rs1990622) causes increased TDP-43 proteinopathy. Specifically, we monitored the development of cytoplasmic TDP-43 aggregates in live cells 24, 48, and 72 hours after knockdown with shRNA targeting *TMEM106B* vs. a scrambled shRNA control.

As shown in Supplementary Fig. 5, cells with *TMEM106B* knockdown trended towards increased TDP-43 aggregation at 48 hours, with significant differences in the number of cells with TDP-43 aggregates at 72 hours, comparing *TMEM106B* knockdown cells vs. control. Moreover, by fixing cells, treating with Triton X-100 to remove soluble proteins, and then staining for insoluble TDP-43 72 hours after *TMEM106B* knockdown, we found that *TMEM106B* knockdown resulted in increased numbers of cells with Triton-insoluble TDP-43 aggregates (Fig. 5). Moreover, by staining cells with an antibody specific for TDP-43 phosphorylated at residues 409/410 (ID3 antibody), we confirmed that at least some of the TDP-43 aggregates seen in the GFP-NLS-TDP system were hyperphosphorylated (Supplementary Fig. 7).

**Fig. 5.**
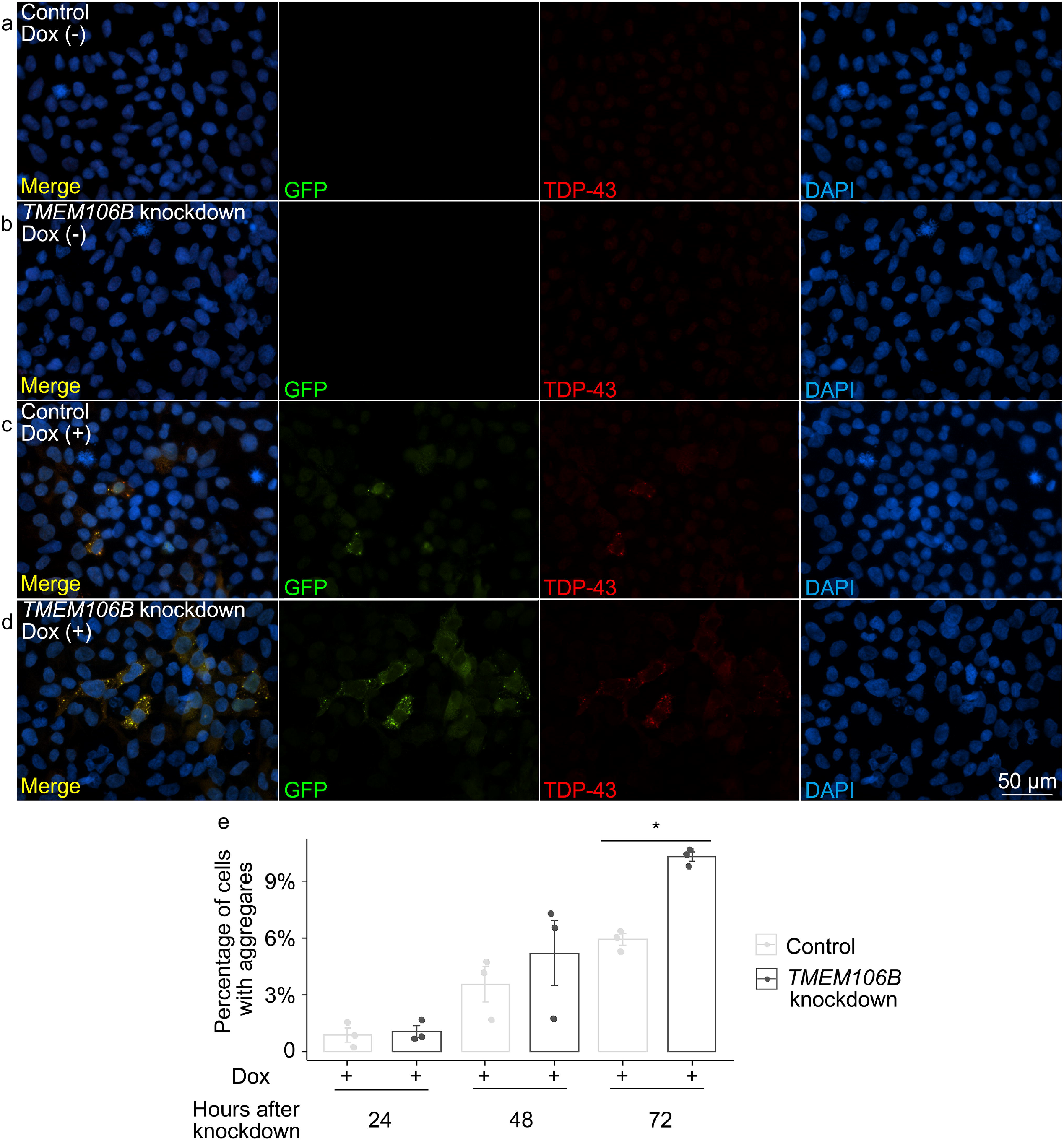
*TMEM106B* knockdown leads to increased propensity for TDP-43 aggregate formation in an inducible cell system that expresses mislocalized TDP-43 protein. **a**-**d** Representative images of double-label immunofluorescence analysis to detect insoluble TDP-43 proteins in iGFP-NLSm cells in the absence (**a**, **b**) or presence (**c**, **d**) of doxycycline after 72 hours of *TMEM106B* knockdown. Triton X-100 was used to extract soluble proteins prior to immunolabeling. Green = GFP, reflecting exogenously-expressed GFP-tagged NLSm TDP-43; red = TDP-43 detected with the C1038 anti-TDP-43 antibody. RIPA-insoluble TDP-43 aggregates were detected after doxycycline induction of iGFP-NLSm cells, with *TMEM106B* knockdown leading to increased numbers of cells with insoluble TDP-43 aggregates. Scale bar = 50 μm. **e** Quantification of cells with TDP-43 aggregates at 48 and 72 hours after *TMEM106B* knockdown (*n* = 3 independent experiments). * One-way ANOVA (*TMEM106B* knockdown vs. control), with Tukey posthoc pairwise comparison p < 0.05 corrected for multiple comparisons.

In order to confirm the shift from soluble to insoluble forms of TDP-43 under conditions of *TMEM106B* knockdown, we performed sequential extractions of total protein lysate 48 and 72 hours after *TMEM106B* knockdown. Immunoblot analysis revealed a 2.5-fold increase in the RIPA-insoluble fraction of endogenous TDP-43 48 hours after *TMEM106B* knockdown (Fig. 6a-f). At 72 hours, the RIPA-insoluble fraction of both endogenous TDP-43 and exogenously-expressed GFP-TDP-43 increased in cells with *TMEM106B* knockdown, compared to controls (Fig. 6g-l).

**Fig. 6.**
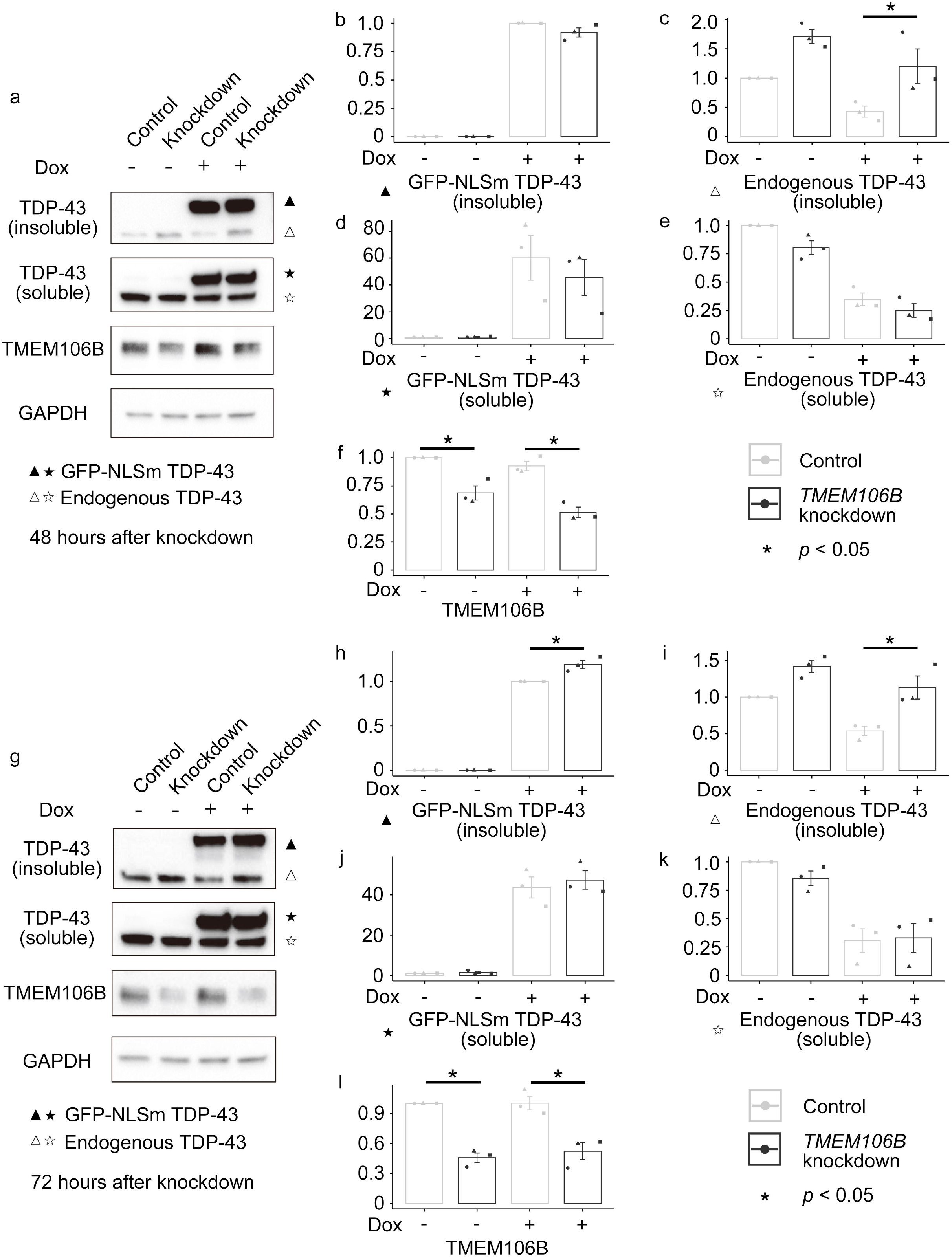
*TMEM106B* knockdown leads to increased insoluble TDP-43 proteins in an inducible cell system that expresses mislocalized TDP-43 protein. Sequential extractions of total protein lysate were performed at 48 and 72 hours after *TMEM106B* knockdown. **a-f** Representative immunoblot pictures of cell lysates at 48 hours after *TMEM106B* knockdown and associated densitometric analysis (*n* = 3 independent experiments) reveal a 2.5-fold increase in the RIPA-insoluble fraction of endogenous TDP-43 (**a, c**). **g-l** Representative immunoblot pictures of cell lysates at 72 hours after *TMEM106B* knockdown and associated densitometric analysis (*n* = 3 independent experiments) revealed a significant increase in the RIPA-insoluble fraction of both endogenous TDP-43 (**g**, **i**) and exogenously-expressed GFP-TDP-43 (**g**, **j**). * One-way ANOVA (*TMEM106B* knockdown vs. control), with Tukey posthoc pairwise comparison p < 0.05 corrected for multiple comparisons.

We have previously shown that increased expression of *TMEM106B* results in grossly abnormal lysosomal appearance and impaired lysosomal acidification [4,2]. *TMEM106B* knockdown, in contrast, has been reported to result in relatively subtle defects in lysosomal positioning [18]. Because altered lysosomal function might result in increased aggregation of TDP-43, we investigated the effects of *TMEM106B* knockdown on lysosomal appearance and acidification in the GFP-NLS-TDP system.

Knockdown of *TMEM106B* did not result in major changes in lysosomal acidification, as assessed using pH sensitive dyes (Supplementary Fig. 6a-6d). Moreover, knockdown of *TMEM106B* in the GFP-NLS-TDP system did not substantially alter the size or appearance of LAMP1-positive organelles (Supplementary Fig. 6e-6h).

Taken together, work in this cell-based system supports a causal role for *TMEM106B* in modifying the development of insoluble TDP-43 aggregates, in the absence of major lysosomal abnormalities.

### TDP-43 pathology and cognitive performance

Our neuropathological studies and cell-based experiments both support the hypothesis that *TMEM106B* genotypes, and by extension *TMEM106B* expression levels, impact TDP-43 proteinopathy in ALS. Intriguingly, however, we previously found that the rs1990622 genotype linked to *decreased* risk for developing FTLD-TDP [4,23] (and lower *TMEM106B* expression levels [8]) were here associated with *more* TDP-43 pathology in the ALS postmortem cases. Knockdown of *TMEM106B* in cells similarly led to more insoluble TDP-43 aggregates. As clinical symptomatology might, in the end, be the most important phenotype, we asked whether the ALS cases with higher global TDP-43 pathology scores showed more or less cognitive impairment.

Out of the 110 ALS autopsy cases, 52 had antemortem cognitive data available for a test of letter-guided verbal fluency (FAS test). In these individuals, higher global TDP-43 pathology scores associated with poorer performance (*β* =−3.228, *p* = 0.029). *TMEM106B* genotypes were not significantly associated with cognitive performance in this limited sample set. In a multivariate linear regression model predicting cognitive score based on *TMEM106B* genotypes, covarying for age at testing, disease duration at testing, sex, TDP-43 pathology score, and the interaction between *TMEM106B* genotype and disease duration, only TDP-43 pathology score associated significantly with cognitive score.

## DISCUSSION

In this study, we investigated candidate genetic modifiers of TDP-43 pathology in a Penn-based autopsy cohort of 899 cases encompassing a spectrum of neurodegenerative diseases. We demonstrate that *TMEM106B* genotypes linked to FTLD-TDP risk associate with the degree of TDP-43 pathology found in ALS, but not in FTLD-TDP, AD, or LBD. We further show that *C9orf72* expansions associate with the degree of TDP-43 pathology found in ALS and FTLD-TDP. Using an inducible cell system that expresses mislocalized TDP-43 protein, we find that knockdown of *TMEM106B* leads to increased TDP-43 aggregation. Finally, in ALS cases with antemortem cognitive data available, we show that higher global TDP-43 pathology scores associate with poorer cognition. Together, the clinical, genetic, neuropathological, and cell-based model data presented in the current study support a model whereby *TMEM106B* plays a causal role in modifying the development of TDP-43 pathology (Fig. 7).

**Fig. 7.**
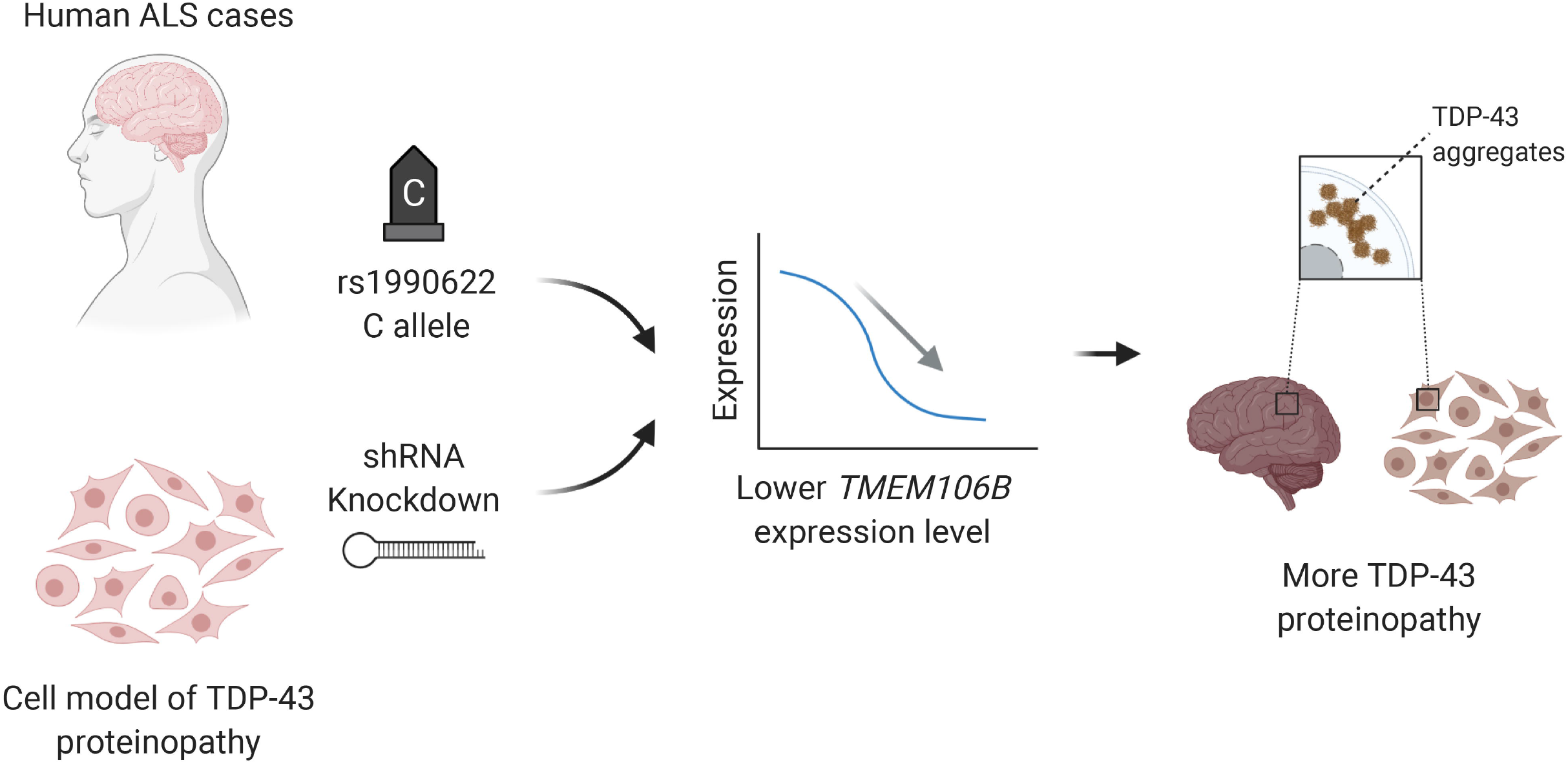
Model of causal effects of *TMEM106B* modulating TDP-43 pathology in ALS. (1) Carrying the minor (C) allele at rs1990622 or (2) *TMEM106B* knockdown in cell models both lead to lower *TMEM106B* gene expression levels. Lower *TMEM106B* expression, in turn, results in development of more TDP-43 pathology.

In our ALS neuropathological studies, the C allele at rs1990622 (associated with lower *TMEM106B* gene expression [8]) was linked to more TDP-43 pathology, reflected both in global TDP-43 pathology scores and in pathology scores for individual CNS regions. Our finding is unexpected since carriers of the C allele at rs1990622 were less likely to develop FTLD-TDP in our original GWAS linking *TMEM106B* to the disease [23], less likely to develop cognitive decline in PD [22], and less likely to have cognitive impairment in our 2011 report of 32 ALS cases with genetic and cognitive data [24]. While surprising, several points add confidence to our finding. First, our sample size of 110 extensively-characterized ALS neuropathology cases is relatively large. Second, our AIM analysis decreases the possibility that differences in genetic ancestry are driving our results. Third, findings from our cell-based model echo the neuropathological results, since decreasing *TMEM106B* expression levels by 30-50% via shRNA-mediated knockdown led to the formation of more TDP-43 aggregates. Finally, a recent *in vivo* study also reported that knockout of *TMEM106B* in mice led to significant increase in levels of insoluble, hyperphosphorylated TDP-43 in the brain [7]. Thus, multiple lines of evidence support the idea that *TMEM106B* is upstream of TDP-43, and *TMEM106B* expression levels modify the development of TDP-43 pathology. While the precise mechanism by which this occurs is unknown, prior work supporting a role for TMEM106B in the function and distribution of lysosomes [1,4,8] suggests a biological pathway worthy of future investigation. We note, however, that nuanced experiments will be needed, as lysosomes do not show gross abnormalities upon *TMEM106B* knockdown in the GFP-NLS-TDP system used here.

A key strength of our study is the combination of detailed, large-scale correlational studies in patient-derived samples with bench-based manipulations in a model system. The failure, to date, of many cell and mouse model-based findings to translate to clinical utility in neurodegenerative disease underscores the importance of verifying that biological pathways and molecular players show signal in human clinical cases: our use of 899 human disease cases with in-depth neuropathological characterization grounds this study in clinical relevance. Clinical studies alone, however, frequently cannot assign directionality of cause and effect. Our direct manipulation of *TMEM106B* levels in a cell-based model allows us to address this gap.

Limitations of this study should be acknowledged along with its strengths. First, the cell-based model used here is non-neuronal, and our findings may not extend to all cell types. Moreover, the aggregates seen in this model may represent pathological forms of TDP-43, stress granules, or other cell biological phenomena, and future experiments exploring these possibilities would add to our understanding. We chose to employ this system because TDP-43 aggregates do not readily form in most neuronal models, and our goal was to determine whether TMEM106B knockdown was sufficient to modify the development of TDP-43 aggregates, requiring a system in which low levels of TDP-43 aggregates occur. Second, cognitive data was available for only about half of the ALS neuropathology cohort, limiting interpretation of the cognitive consequences of *TMEM106B* genotype and TDP-43 pathology. In this regard, studies in ALS with larger sample sizes and a wider range of cognitive tests would be a valuable extension to the work presented here.

Limitations notwithstanding, our study sheds new light on the relationship between TMEM106B and TDP-43, and, in particular, supports a causal role for *TMEM106B* expression levels in modifying the development of TDP-43 proteinopathy. Additionally, it exemplifies an approach combining large-scale characterization of patient-derived tissues with hypothesis-driven bench manipulation. Given the large number of ALS patients with no proven neuroprotective therapies, our demonstration that TMEM106B may play a disease-modifying role paves the way for work investigating its potential as a therapeutic target.

## ACKNOWLEDGMENTS

We thank our patients and their families for contributing to this research.

This study was supported by the NIH (RO1 NS082265, P30 AG010124, U19 AG062418). Fei Mao is additionally supported by the National Natural Science Foundation of China (81700053) and Alice Chen-Plotkin by the Chan Zuckerberg Initiative Neurodegeneration Challenge, the American Heart Association/Allen Institute Brain Health Initiative, and the Parker Family Chair.

