## Supplementary materials for "TMEM106B modifies TDP-43 pathology in human ALS brain and cell-based models of TDP-43 proteinopathy"

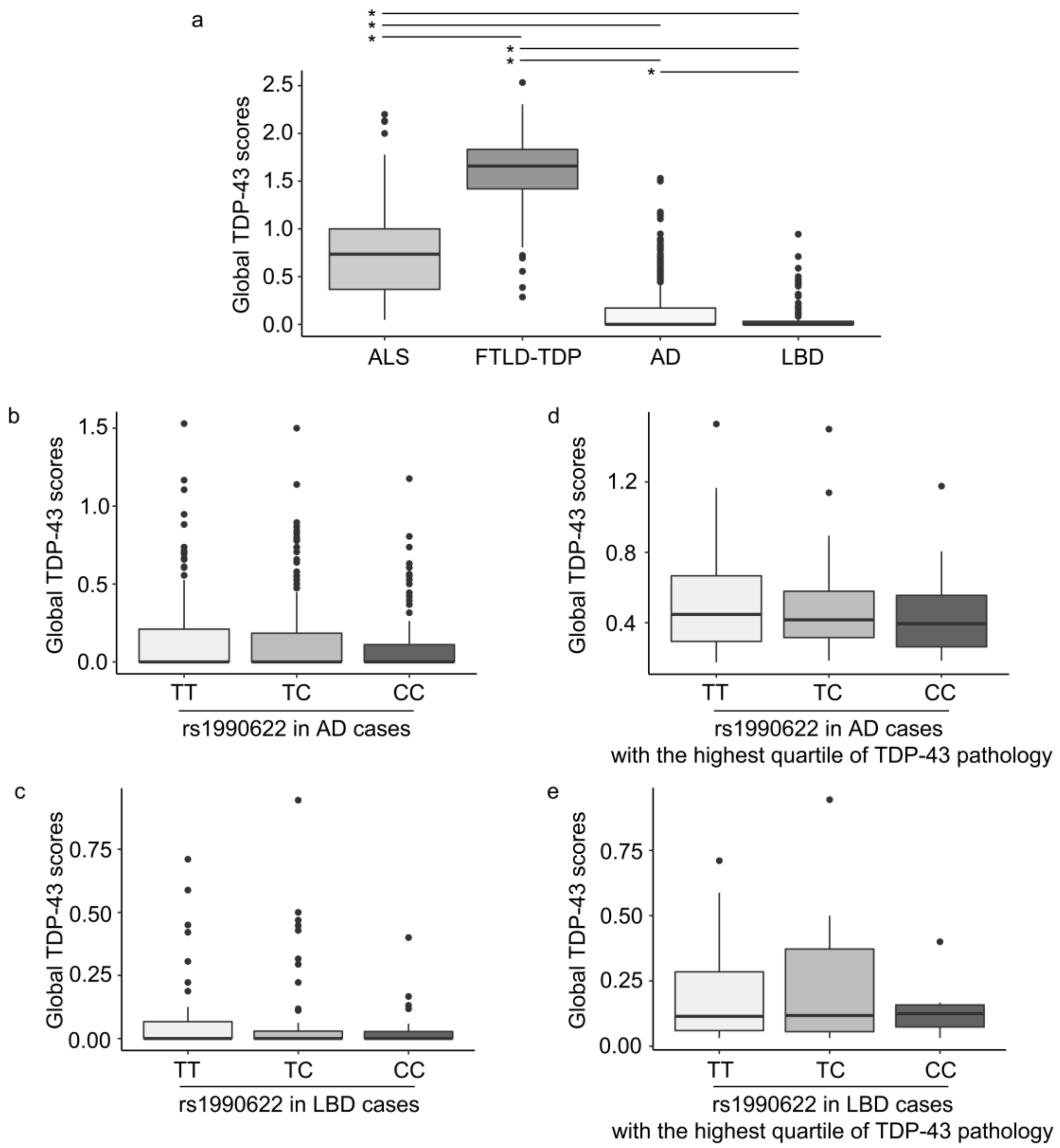


**Supplementary Fig. 1**

Boxplots comparing TDP-43 pathology score among neurodegenerative diseases. FTLD-TDP cases had the highest global TDP-43 scores, with a median score of 1.66 (IQR = 1.41-1.83), followed by ALS cases (median score 0.74, IQR 0.37-1.00). AD and LBD cases showed low levels of TDP-43 pathology (**a**). All groups differ from each other significantly in degree of TDP-43 pathology observed, $p < 0.001$ for ALS and FTLD, $p < 0.001$ for ALS and AD, $p < 0.001$ for ALS and LBD, $p < 0.001$ for FTLD and AD, $p < 0.001$ for FTLD and LBD, $p = 0.012$ for AD and LBD (**a**). Global TDP-43 pathology scores for AD cases (**b**) and LBD cases (**c**) by *TMEM106B* genotype.

Global TDP-43 pathology scores for cases with the highest quartile of TDP-43 pathology for AD cases (**d**) and LBD cases (**e**) by *TMEM106B* genotype. (* Statistically significant difference)

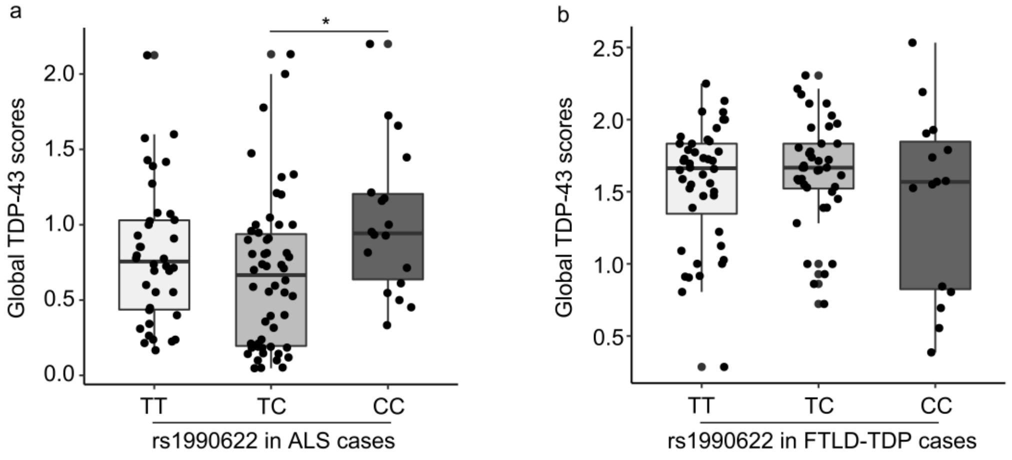


**Supplementary Fig. 2**

Boxplots comparing TDP-43 pathology scores among *TMEM106B* rs1990622 genotypes (TT, TC and CC) in ALS and FTLD-TDP. **a.** For ALS cases, global TDP-43 pathology scores differed significantly among *TMEM106B* genotypes, with homozygotes for the minor allele (CC) having the highest scores (* $p$ = 0.020). **b.** Global TDP-43 scores did not differ among carriers of different *TMEM106B* genotypes in FTLD-TDP cases.

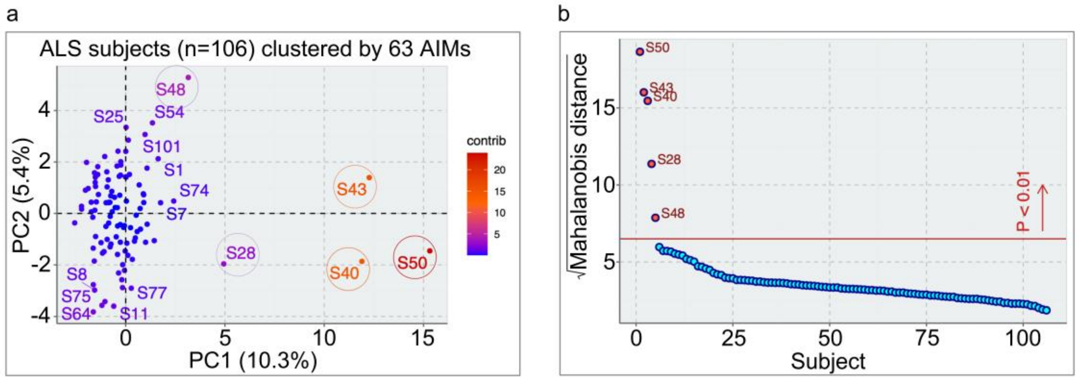


**Supplementary Fig. 3**

We analyzed our ALS cohort for individuals with significantly different genetic ancestry using 63 Ancestry Informative Markers (AIMs). **a**. Principal component analysis (PCA) of 106 ALS cohort patients was performed using 63 AIMs. Clustering of ALS cohort patients by the first two principal components (each dot represents one subject, colored according to contribution to the principal components) shows three well-separated subjects (S40, S43, S50) along the first principal component (PC1), representing 10.3% of the total variance. S28 and S48 are separated, to a lesser extent, along the first and second principal components. **b**. Five outlier individuals (red dots, $p < 0.01$) with significantly different genetic ancestry were detected by multivariate distance analysis (Mahalanobis distance) of the first 10 PCs simultaneously. P-values were estimated by cumulative probability of the chi-squared distribution and adjusted for multiple hypothesis testing by Bonferroni correction.

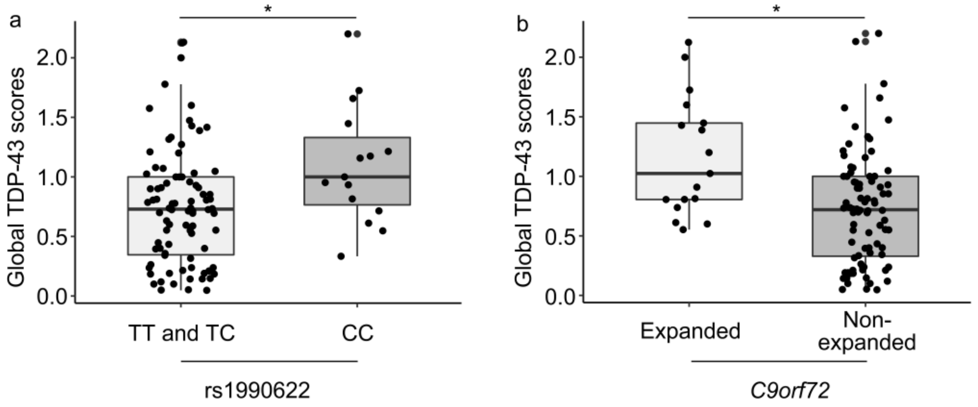


c Multivariate model: TDP-43 score predicted by *TMEM106B* genotype, *C9orf72* expansion and age at death.

| | $R^2$ | Coefficient | *p* value |
|---|---|---|---|
| Overall model | 0.171 | | |
| *TMEM106B* genotype (major allele) | | -0.330 | 0.012* |
| *C9orf72* expansion | | 0.429 | 0.001* |
| Age at death | | -0.009 | 0.058 |

* Statistically significant

**Supplementary Fig. 4**

In the subgroup of 101 individuals remaining in our ALS cohort after elimination of cases with significantly different genetic ancestry, *TMEM106B* genotype and presence of *C9orf72* expansion both continue to associate with the global extent of TDP-43 pathology. **a**. Global TDP-43 pathology scores differed significantly among *TMEM106B* genotypes under the major allele (T) dominant model ($p$ = 0.011). **b**. *C9orf72* expansion carriers showed higher global TDP-43 pathology scores ($p$ = 0.002). c. Excluding five individuals of different genetic ancestry, as well as the four individuals for whom we could not ascertain AIMs, neither the association between *TMEM106B* genotype and global TDP-43 pathology scores, nor the association between *C9orf72* expansion status and global TDP-43 pathology scores were substantially affected in a multivariate linear regression predicting TDP-43 pathology score using *TMEM106B* genotype, *C9orf72* expansion status, and age at death.

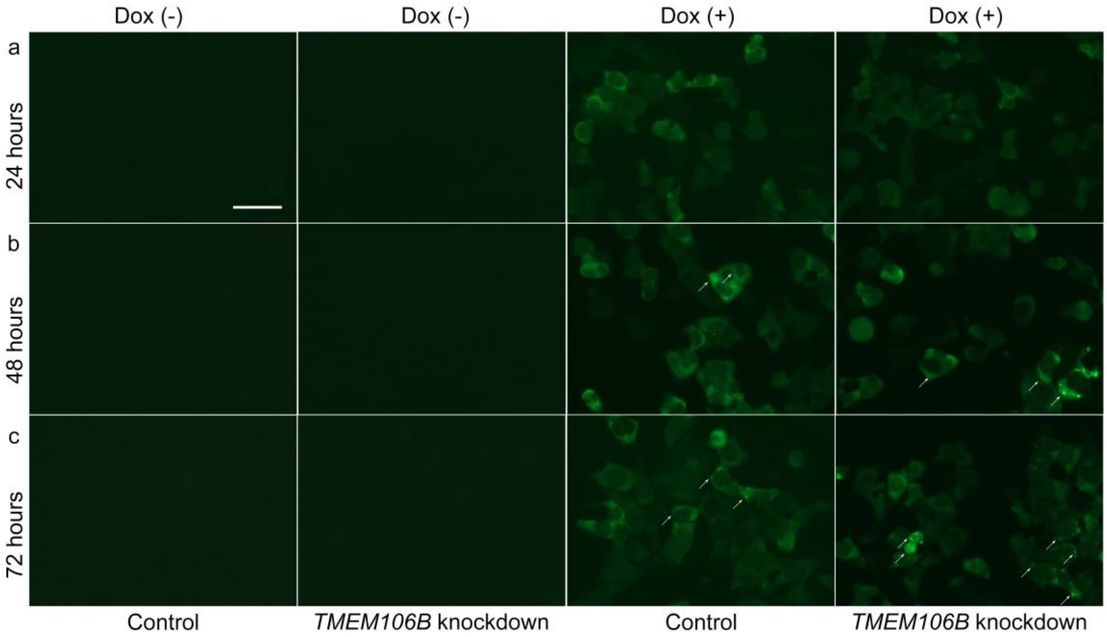


**Supplementary Fig. 5**

*TMEM106B* knockdown leads to more cells with TDP-43 aggregates in cell-based models of TDP-43 proteinopathy. Representative photomicrographs of live cells at 24 hours (**a**), 48 hours (**b**) and 72 hours (**c**) after *TMEM106B* knockdown. Expression of exogenous doxycycline (Dox)-inducible GFP-TDP-43 proteins (green) was detected in the presence of Dox (right 2 columns). Compared with control, *TMEM106B* knockdown cells trended towards increased numbers of cells with TDP-43 aggregation (arrows) at 48 hours, with significant differences in the number of cells with TDP-43 aggregates (arrows) at 72 hours. Scale bar = 50 µm.

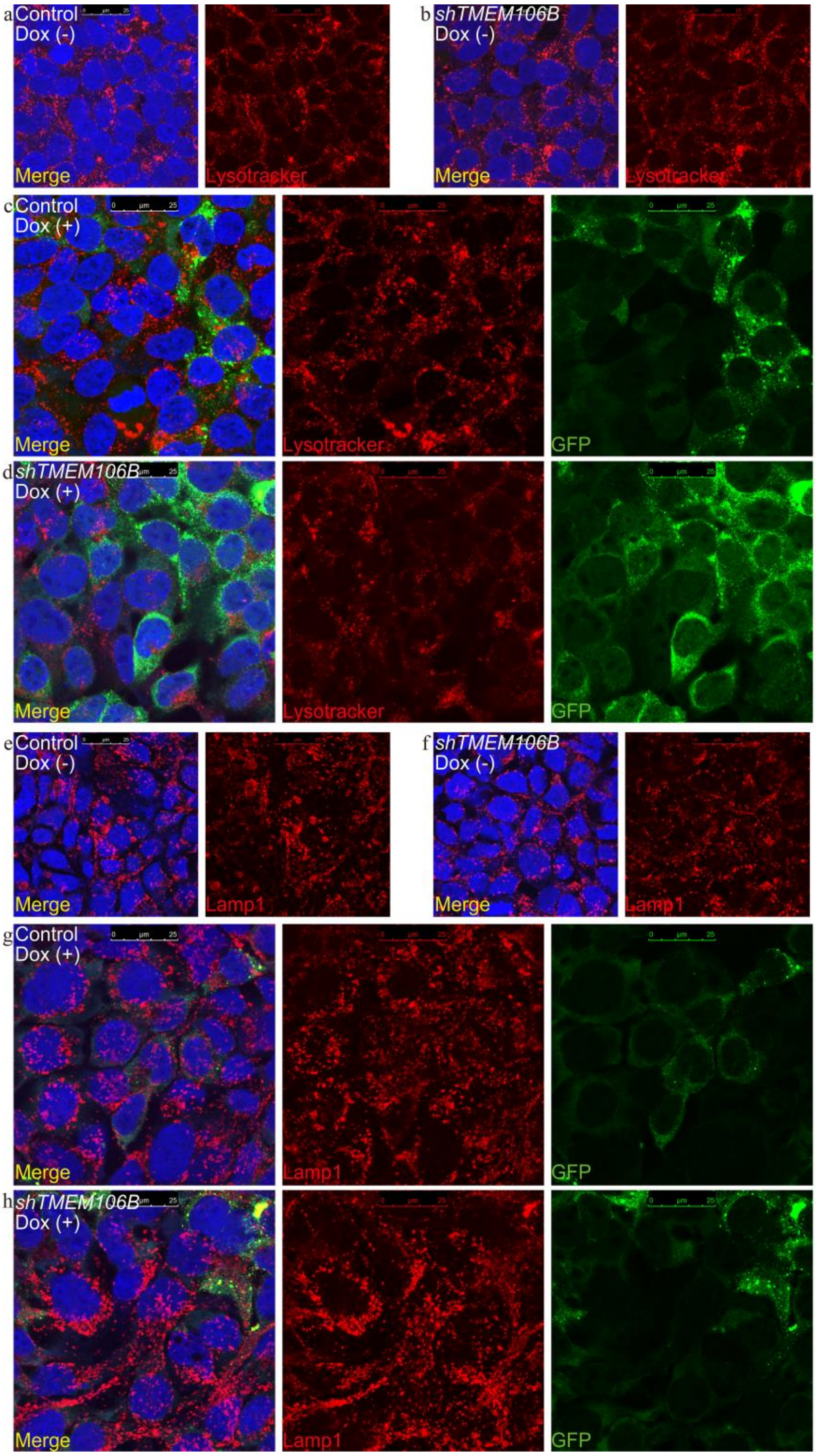


**Supplementary Fig. 6**

*TMEM106B* knockdown does not lead to a change in lysosomal morphology. Representative images of cells stained with the pH-sensitive dye Lysotracker (in red, panels **a-d**), without (**a-b**) or with (**c-d**) GFP-TDP-43 induction under control shRNA (**a**, **c**) or *TMEM106B* knockdown (**b**, **d**) conditions are shown. Lamp1 staining with the H4A3 antibody (in red, panels **e-h**) also shows a lack of maor morphological lysosomal changes without (**e-f**) or with (**g-h**) GFP-TDP-43 induction under control shRNA (**e**, **g**) or *TMEM106B* knockdown (**f**, **h**) conditions.

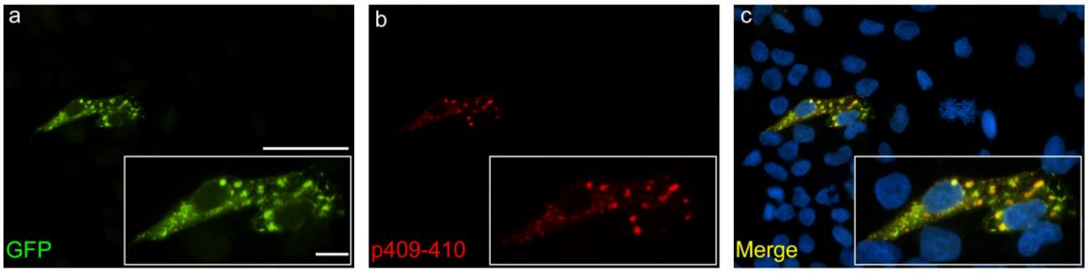


**Supplementary Fig. 7**

A subset of the TDP-43 aggregates in iGFP-NLSm cells are phosphorylated. Representative double-label IF images of GFP (**a**) and p409-410 (**b**). p409-410 TDP-43 (stained with ID3 antibody) partially co-localizes with the GFP-TDP-43 NLSm protein (**c**), indicating the aggregates in iGFP-NLSm cells are a mix of phosphorylated and non-phosphorylated TDP-43 protein. Insets are higher magnifications of the cells. Scale bar = 50 µm and 10 µm (insets).

**Supplementary Table 1.** Ancestry Informative Markers

| | RsID | Chromosome | Position | Alleles |
|---|---|---|---|---|
| 1 | rs2986742 | 1 | 6550376 | T/C |
| 2 | rs4908343 | 1 | 27931698 | G/A |
| 3 | rs3118378 | 1 | 68849687 | A/G |
| 4 | rs3737576 | 1 | 101709563 | T/C |
| 5 | rs7554936 | 1 | 151122489 | C/T |
| 6 | rs1407434 | 1 | 186149032 | G/A |
| 7 | rs798443 | 2 | 7968275 | G/A |
| 8 | rs7421394 | 2 | 14756349 | A/G |
| 9 | rs4666200 | 2 | 29538411 | G/A |
| 10 | rs13400937 | 2 | 79864923 | T/G |
| 11 | rs260690 | 2 | 109579738 | C/A |
| 12 | rs1569175 | 2 | 201021954 | T/C |
| 13 | rs9809104 | 3 | 39146429 | T/C |
| 14 | rs6548616 | 3 | 79399575 | T/C |
| 15 | rs12629908 | 3 | 120522716 | G/A |
| 16 | rs9845457 | 3 | 135914476 | G/A |
| 17 | rs1513181 | 3 | 188574996 | G/A |
| 18 | rs9291090 | 4 | 5390637 | A/C |
| 19 | rs10007810 | 4 | 41554364 | G/A |
| 20 | rs7657799 | 4 | 105375423 | T/G |
| 21 | rs2702414 | 4 | 179399523 | G/A |
| 22 | rs37369 | 5 | 35037115 | C/T |
| 23 | rs6451722 | 5 | 43711378 | G/A |
| 24 | rs12657828 | 5 | 79085726 | A/G |
| 25 | rs6556352 | 5 | 155471714 | C/T |
| 26 | rs1040045 | 6 | 4747159 | G/A |
| 27 | rs2397060 | 6 | 51611470 | T/C |
| 28 | rs192655 | 6 | 90518278 | G/A |
| 29 | rs4458655 | 6 | 163221792 | T/C |
| 30 | rs1871428 | 6 | 168665760 | G/A |
| 31 | rs731257 | 7 | 12669251 | G/A |
| 32 | rs7803075 | 7 | 130742066 | A/G |
| 33 | rs6464211 | 7 | 151873853 | C/T |
| 34 | rs1471939 | 8 | 28941305 | C/T |
| 35 | rs10511828 | 9 | 28628500 | T/C |
| 36 | rs2306040 | 9 | 93641199 | T/C |
| 37 | rs4746136 | 10 | 75300994 | G/A |
| 38 | rs4918842 | 10 | 115316812 | T/C |
| 39 | rs11227699 | 11 | 66898492 | G/A |
| 40 | rs2416791 | 12 | 11701488 | A/G |
| 41 | rs772262 | 12 | 56163734 | G/A |

| | | | | |
|---|---|---|---|---|
| 42 | rs9319336 | 13 | 27624356 | T/C |
| 43 | rs7997709 | 13 | 34847737 | C/T |
| 44 | rs9530435 | 13 | 75993887 | T/C |
| 45 | rs9522149 | 13 | 111827167 | T/C |
| 46 | rs1760921 | 14 | 20818131 | T/C |
| 47 | rs8021730 | 14 | 67886781 | G/T |
| 48 | rs946918 | 14 | 83472868 | G/T |
| 49 | rs200354 | 14 | 99375321 | G/T |
| 50 | rs3784230 | 14 | 105679055 | A/G |
| 51 | rs8035124 | 15 | 92105708 | A/C |
| 52 | rs4984913 | 16 | 740466 | A/G |
| 53 | rs10512572 | 17 | 69512099 | G/A |
| 54 | rs4798812 | 18 | 9420504 | G/A |
| 55 | rs881728 | 18 | 59333108 | C/A |
| 56 | rs4891825 | 18 | 67867663 | G/A |
| 57 | rs874299 | 18 | 75056284 | T/C |
| 58 | rs8113143 | 19 | 33652247 | C/A |
| 59 | rs3745099 | 19 | 52901905 | G/A |
| 60 | rs2532060 | 19 | 55614923 | T/C |
| 61 | rs6104567 | 20 | 10195433 | T/G |
| 62 | rs3907047 | 20 | 54000914 | T/C |
| 63 | rs4821004 | 22 | 32366359 | C/T |

**Supplementary Table 2.** Bivariate linear model: TDP-43 scores predicted by *TMEM106B* genotype in FTLD-TDP

| | Codominant model | | Major allele (T) dominant model | |
|---|---|---|---|---|
| | Coefficient | *p* value | Coefficient | *p* value |
| Amygdala | 0.077 | 0.537 | -0.036 | 0.886 |
| Angular gyrus | -0.103 | 0.403 | 0.325 | 0.178 |
| Anterior cingulate | -0.065 | 0.492 | 0.209 | 0.262 |
| Cerebellum | 0.027 | 0.492 | -0.116 | 0.138 |
| Caudate/Putamen | -0.122 | 0.364 | 0.555 | 0.038* |
| CA1/subiculum | 0.103 | 0.406 | 0.022 | 0.929 |
| Dentate gyrus | -0.049 | 0.824 | 0.773 | 0.086 |
| Entorhinal cortex | -0.031 | 0.755 | 0.211 | 0.288 |
| Globus Pallidus | 0.056 | 0.692 | 0.187 | 0.504 |
| Locus coeruleus | -0.063 | 0.466 | 0.050 | 0.768 |
| Motor cortex | -0.066 | 0.717 | 0.191 | 0.584 |
| Midbrain | -0.063 | 0.637 | 0.197 | 0.464 |
| Medulla | 0.005 | 0.97 | 0.074 | 0.781 |
| Middle frontal cortex | -0.146 | 0.204 | 0.565 | 0.013* |
| Occipital cortex | 0.020 | 0.885 | -0.115 | 0.671 |
| Pons | 0.028 | 0.800 | 0.081 | 0.709 |
| Spinal cord | 0.021 | 0.924 | 0.088 | 0.832 |
| Superior/middle temporal cortex | 0.055 | 0.545 | -0.027 | 0.881 |
| Substantia nigra | -0.053 | 0.655 | 0.025 | 0.915 |
| Thalamus/subthalamus | -0.084 | 0.539 | 0.090 | 0.748 |

* Nominal $p < 0.05$

**Supplementary Table 3.** Bivariate linear model: TDP-43 scores predicted by *C9orf72* expansion in FTLD-TDP

| | *C9orf72* expansion | |
|---|---|---|
| | Coefficient | *p* value |
| Amygdala | 0.351 | 0.092 |
| Angular gyrus | 0.154 | 0.462 |
| Anterior cingulate | 0.186 | 0.241 |
| Cerebellum | 0.037 | 0.578 |
| Caudate/Putamen | 0.083 | 0.714 |
| CA1/subiculum | 0.103 | 0.630 |
| Dentate gyrus | 0.922 | 0.005** |
| Entorhinal cortex | 0.233 | 0.174 |
| Globus Pallidus | 0.043 | 0.856 |
| Locus ceruleus | 0.095 | 0.519 |
| Motor cortex | 0.302 | 0.384 |
| Midbrain | 0.571 | 0.009** |
| Medulla | 0.432 | 0.054 |
| Middle frontal cortex | 0.398 | 0.039* |
| Occipital cortex | 0.375 | 0.102 |
| Pons | 0.102 | 0.579 |
| Spinal cord | 0.503 | 0.202 |
| Superior/middle temporal cortex | 0.129 | 0.406 |
| Substantia nigra | 0.475 | 0.016* |
| Thalamus/subthalamus | 0.487 | 0.031* |

* Nominal $p < 0.05$, ** Nominal $p < 0.01$
